# *APOE* ε4 defines a systemic immune endophenotype independent of clinical trajectory in amyotrophic lateral sclerosis

**DOI:** 10.64898/2026.02.02.703399

**Authors:** Artur Shvetcov, Shannon Thomson, Sterling Kwan, Terri G. Thompson, Jeffery D. Rothstein, Caitlin A. Finney

**Affiliations:** Neurodegeneration and Disease Modelling Lab, Westmead Institute for Medical Research, The University of Sydney, Westmead, New South Wales, Australia; School of Medical Sciences, Faculty of Medicine and Health, The University of Sydney, Sydney, New South Wales, Australia; OnPoint Scientific, Inc., San Diego, California, USA; Brain Science Institute, Johns Hopkins University School of Medicine, Baltimore, Maryland, USA; Department of Neurology, Johns Hopkins University School of Medicine, Baltimore, Maryland, USA

**Author notes:** Corresponding authors: E; E; A: 176 Hawkesbury Road, Westmead, NSW, Australia 2145. These authors contributed equally (Artur Shvetcov, Caitlin A. Finney).

**Keywords:** Amyotrophic lateral sclerosis, multiomics, APOE ε4, motor neurons, plasma, immune

## Abstract

**Background:** Amyotrophic lateral sclerosis (ALS) is clinically heterogeneous, and genetic modifiers may drive molecular endophenotypes without obvious clinical stratification. The apolipoprotein E ε4 (*APOE* ε4) allele is a major Alzheimer’s disease risk allele, but its biological impact in ALS remains unclear.

**Methods:** Using the Answer ALS cohort, longitudinal motor, cognitive, and neuropsychiatric measures were modelled using mixed-effects approaches. Patient induced pluripotent stem cell-derived motor neuron multiomics (chromatin accessibility, transcriptomics, and proteomics) were analysed using supervised machine learning. Plasma SomaScan profiling was used to derive an *APOE* ε4-associated protein signature and to test its stability across serial visits, biological pathway enrichment, and associations with clinical progression.

**Results:** *APOE* ε4 carriage was not associated with baseline severity or rate of functional decline and showed no consistent effects on cognitive or neuropsychiatric trajectories. Motor neuron multiomic profiles similarly demonstrated no reproducible *APOE* ε4 signal and did not reliably classify genotype. In contrast, plasma proteomics identified an *APOE* ε4 protein signature that classified ε4 status with high accuracy in ALS (AUC 0.98) and non-ALS motor neuron disease (AUC 0.86) and was enriched for immune and inflammatory biology. This systemic signature was highly stable across repeated sampling, indicating a persistent genotype-associated state. Within this plasma endophenotype, a small set of proteins tracked functional decline and a composite score stratified fast versus slow progression. Baseline composite scores were elevated in *APOE* ε4 carriers in both ALS and neurologically unimpaired controls, consistent with a stable systemic shift detectable beyond overt disease status.

**Conclusions:** *APOE* ε4 defines a persistent, immune-enriched systemic proteomic endophenotype in ALS that is not captured by clinical trajectories or motor neuron-only profiling yet relates to disease progression. Plasma-based, genotype-informed endophenotyping offers a translational pathway for biomarker stratification and therapeutic prioritisation in ALS and potentially other heterogeneous neurodegenerative disorders.

## Background

Amyotrophic lateral sclerosis (ALS) is a fatal neurodegenerative disease characterized by progressive motor neuron loss and substantial clinical heterogeneity (1, 2). Despite growing interest in the genetic modifiers of ALS, the role of the apolipoprotein E ε4 (*APOE* ε4) allele, a major risk factor for Alzheimer’s disease (AD), remains controversial. Several large-scale studies have reported no association between *APOE* ε4 and ALS risk, age of onset, or progression (3, 4, 5, 6, 7, 8, 9, 10, 11). Even in studies that have identified higher *APOE* ε4 frequency (10) or earlier onset in subgroups like bulbar onset ALS (12), no consistent genotype-phenotype pattern has emerged. Genotype relevance is often interpreted through the presence or absence of overt clinical stratification and therefore these findings have fostered the widespread assumption that *APOE* ε4 is not functionally relevant in ALS pathogenesis (3, 7, 8, 13, 14). However, as seen in other neurodegenerative diseases, clinically indistinguishable subgroups can have profoundly different underlying biology. Parkinson’s disease (PD), for example, carriers of the pathogenic LRRK2 p.R1441C mutation often present with a clinical profile indistinguishable from sporadic PD, despite clear molecular pathogenicity (15). Thus, the absence of a genotype-phenotype correlation should not be interpreted as absence of biological effect. Instead, genotypes may define molecular endophenotypes that shape disease biology without dictating clinical trajectory. Overall, this conceptual misunderstanding has limited the field’s ability to detect subtle, systems-level effects of common variants like *APOE* ε4.

Emerging data highlight this view. In ALS mouse models, *APOE* acts as a key regulator of the microglial state. Here, activation of the TREM2-APOE axis drives a shift from homeostatic to neurodegenerative microglia, while inhibition of this pathway restores microglial function and rescues neurons (16). Further, *APOE* ε4 has been linked to impaired antioxidant function. One study showed that it has a reduced ability to neutralize lipid peroxidation products like 4-hydroxynonenal (4-HNE), leading to increased oxidative stress and vulnerability in mouse motor neurons (17). These mechanisms are directly relevant to ALS, where immune activation and oxidative injury are key pathogenic features (18, 19). In humans, indirect evidence also suggests systemic consequences of *APOE* ε4 in ALS. *APOE* ε4 has been associated with increased cerebral amyloid angiopathy and reduced cerebrospinal fluid (CSF) Aβ_42/40_ levels in ALS patients, hinting at co-pathology or shared pathways with AD (20, 21). In some ALS cohorts, *APOE* ε4 carriers show greater executive dysfunction, though findings are inconsistent and often confounded by overlapping neurodegenerative processes (7, 21). *APOE* ε4 may also interact with environmental exposures. In military veterans, the association between head trauma and ALS risk is significantly stronger among *APOE* ε4 carriers, suggesting gene-environment interactions (22). Despite these insights, systematic multiomic evaluation of the functional impact of *APOE* ε4 in ALS has not been studied.

Here, we address this gap using multiomic and longitudinal data from the Answer ALS cohort, one of the most comprehensive patient-derived datasets in ALS. We first assess whether *APOE* ε4 status influences clinical outcomes, including ALS Functional Rating Scale-Revised (ALSFRS-R) decline, cognitive impairment, and pseudobulbar affect, and confirm previous reports of no detectable phenotype-level effect. We next examined the epigenetic, transcriptomic, and proteomic profiles of patient induced pluripotent stem cell-derived motor neurons (iMNs), revealing no effect of *APOE* ε4 on chromatin accessibility, gene expression, or protein abundance. In stark contrast, plasma proteomic profiling revealed a distinct, reproducible *APOE* ε4-associated signature in ALS patients, marked by robust enrichment of immune and inflammatory pathways. This *APOE* ε4 systemic signature was stable over time, indicating a persistent immune state that precedes and accompanies disease progression. Further, we find that specific proteins within this plasma *APOE* ε4 signature are associated with the rate of clinical disease progression. These findings position *APOE* ε4 as playing a biological role in ALS, not through overt clinical or neuronal effects, but via a stable, peripheral immune phenotype that likely primes carriers for neurodegeneration. This work challenges the assumption that the absence of genotype-phenotype correlation implies biological neutrality and underscores the limitations of conventional approaches that rely on clinical stratification alone. More broadly, these results support a precision medicine framework grounded in molecular endophenotypes rather than clinical presentation, offering a blueprint for identifying disease-modifying processes and therapeutic targets in genetically defined subgroups across heterogeneous neurodegenerative disorders.

## Methods

### Answer ALS cohort

The Answer ALS cohort comprises the largest harmonized collection of longitudinal clinical data and biospecimens from individuals with amyotrophic lateral sclerosis (ALS) assembled to date. Participants were recruited across multiple neuromuscular centers in the US, including Johns Hopkins University, Massachusetts General Hospital, The Ohio State University, Emory University, Washington University, Northwestern University, Cedars-Sinai, and Texas Neurology (23). The present study included individuals with sporadic and familial ALS (ALS), non-ALS motor neuron diseases (non-ALS MND), including primary lateral sclerosis, progressive bulbar palsy, and progressive muscular atrophy, as well as age-matched neurologically unimpaired controls without a personal or family history of ALS (23). Longitudinal clinical progression was assessed over a minimum of three study visits using the ALS Functional Rating Scale-Revised (ALSFRS-R), which captures functional impairment across bulbar, limb, and respiratory domains (23, 24). Pseudobulbar affect was evaluated using the Center for Neurologic Study-Lability Scale (CNS-LS), a seven-item self-report measure of pathological laughing and crying (25, 26). Cognitive impairment was measured using the ALS Cognitive Behavioural Screen (ALS-CBS) (27). Peripheral blood mononuclear cells (PBMCs) were isolated from baseline whole blood samples, and whole-genome sequencing (WGS) was performed to identify ALS-associated genetic variants, including *C9orf72, SOD1, FUS, TARDBP*, and *TBK1* (23). *APOE* genotype was determined by extracting genotypes at the two canonical *APOE*-defining single nucleotide polymorphisms, rs429358 and rs7412, located on chromosome 19. Variant genotypes were queried from compressed VCF files using ‘bcftools’ and exported in tabular format. Genotypes were parsed and harmonised across variants to ensure consistent sample ordering. *APOE* alleles were assigned based on the combination of rs429358 and rs7412 genotypes: ε2 (rs429358 = 0/0, rs7412 = 1/1), ε3 (rs429358 = 0/0, rs7412 = 0/0), and ε4 (rs429358 = 1/1 or 0/1 with rs7412 = 0/0). Individuals carrying at least one ε4 allele (ε2/ε4, ε3/ε4, or ε4/ε4) were classified as *APOE* ε4 carriers, while all others were classified as non-carriers. Genotype calls were linked to participant identifiers using cohort metadata and merged with demographic and clinical information, including age, sex, and diagnostic group. *APOE* ε4 carrier status was used as a binary exposure variable in all downstream analyses. All participants provided written informed consent, and study procedures were approved by the institutional review boards of each participating site.

### Patient iPSC-derived motor neurons and multiomics

Patient induced pluripotent stem cells (iPSCs) were generated from PBMCs using non-integrating episomal factors (23). iPSCs were then differentiated into motor neurons (iMNs) using a staged neural induction, motor neuron specification, and terminal maturation protocol, as described elsewhere (23, 28). Chromatin accessibility, gene expression and protein abundance in iMNs were analysed using ATAC-seq, bulk RNA-seq, and label-free quantitative proteomics, respectively, on day 32 (23). For ATAC-seq and bulk RNA-seq analyses were performed on feature level count matrices. Features with zero total counts, zero variance, or low detection (retaining features with counts >1 in ≥ 20% of samples) were removed prior to downstream analysis. Counts were variance stabilized using ‘DESeq2’ (29). Proteomic intensities were log_2_-transformed following addition of a small offset to accommodate zero values. For each omic dataset, batch-associated variation was adjusted using limma, with sequencing batch specified as the batch factor and a design matrix including diagnosis, sex, and age (30).

### Plasma proteomics

Plasma proteomic profiling was performed using the SomaScan v4.1 platform (SomaLogic), which quantifies approximately 7,000 circulating proteins using an aptamer-based affinity proteomics approach. This technology employs slow off-rate modified aptamers (SOMAmers) containing chemically modified nucleotides to enable high-affinity and high-specificity protein binding (31). Proteomic data generated by SomaLogic were processed using the Adaptive Normalization by Maximum Likelihood (ANML) pipeline, which performs standardization, normalization, and calibration across samples, with final protein abundances reported as relative fluorescent units (RFU). Plasma proteomic profiles were analysed using measurements from baseline samples. Protein identifiers were mapped to UniProt IDs, and samples were annotated with clinical and demographic metadata.

### Feature selection and machine learning

Differential chromatin accessibility, gene expression, and protein abundance associated with *APOE* ε4 carrier status were assessed using linear models with empirical Bayes moderation implemented in limma, modelling *APOE* ε4 status while adjusting for diagnosis, sex, and age, with multiple testing controlled using the Benjamini-Hochberg false discovery rate (FDR). To assess whether *APOE* ε4 status could be captured by coordinated multivariate molecular patterns not detected by univariate analyses, we performed exploratory supervised machine-learning analyses on ATAC-seq, bulk RNA-sequencing, and label-free proteomic data derived from iMNs. Variance stabilized and batch-adjusted feature matrices were used as input, with feature selection performed using mutual information calculated on training data only using ‘FSelectorRcpp’. Supervised random forest classifiers were trained using repeated cross-validation (5-fold cross-validation repeated five times) implemented in ‘caret’, with area under the receiver operating characteristic curve (AUC) as the primary performance metric. To mitigate class imbalance, minority-class upsampling was applied within resampling folds, and model performance was evaluated on held-out test sets.

To identify *APOE* ε4-associated plasma proteins independent of disease effects, mutual information-based feature selection using ‘FSelectorRcpp’ was performed in healthy control participants, ranking proteins by their association with *APOE* ε4 carrier status.

Proteins with non-zero mutual information were retained as candidate features. To evaluate the generalisability of the derived protein set, random forest classifiers were trained using healthy control samples and subsequently applied, without retraining, to ALS and non-ALS motor neuron disease cohorts. Classifier performance was assessed using repeated cross-validation (5-fold cross-validation repeated five times), with AUC calculated from predicted class probabilities as the primary performance metric. To address class imbalance, minority-class upsampling was performed within resampling folds.

### Longitudinal analyses

Longitudinal analyses were performed using regression-based models appropriate to the repeated measures structure of each outcome. Time was defined as years since screening, calculated from the recorded days-since-screening variable divided by 365.25. For clinical outcomes (ALSFRS-R, ALS-CBS, CNS-LS), longitudinal trajectories were analysed using linear mixed-effects models including fixed effects for time, *APOE* ε4 carrier status, and their interaction, with additional adjustment for age and sex. Participant-specific random intercepts and, where supported by the data, random slopes for time were included to capture heterogeneity in baseline severity and change over time. Models were fit by restricted maximum likelihood, with statistical inference for fixed effects obtained using Satterthwaite’s degrees-of-freedom approximation (‘lmerTest’).

To quantify stability of *APOE* ε4-associated plasma effects across visits, *APOE* ε4 effect sizes were estimated separately at each timepoint for each protein using linear models, and concordance across visits was assessed by pairwise correlations of effect estimates. To relate longitudinal molecular change to clinical progression, individual ALSFRS-R decline rates were estimated using mixed-effects models with random slopes, and per-patient protein slopes were derived by regressing log -transformed protein abundance on time. Protein slopes were then regressed on ALSFRS-R slope and *APOE* ε4 status, and FDR was controlled across proteins using the Benjamini-Hochberg procedure. Composite scores were derived by principal component analysis of progression-associated protein slopes, with associations with ALSFRS-R decline assessed by correlation, linear regression, and logistic regression in tertile-defined fast versus slow progressors.

### Enrichment analyses

Functional enrichment analyses of *APOE* ε4 plasma proteins were performed using NetworkAnalyst (v3.2) (32, 33, 34). Protein-protein interaction networks were constructed using a first-order network in the International Molecular Exchange Consortium (IMEx) Interactome (35). Enrichment of biological processes was assessed using the Protein Analysis Through Evolutionary Relationships (PANTHER) classification system (36, 37), while pathway-level enrichment was evaluated using the Kyoto Encyclopedia of Genes and Genomes (KEGG) (38, 39). For all enrichment analyses, statistical significance was reported using FDR-adjusted p values. Protein-drug interaction analyses were also conducted in NetworkAnalyst (v3.2) using curated interactions from the DrugBank database (v 5.1.13) (40, 41).

## Results

### Answer ALS cohort demographics

To clarify the biological relevance of *APOE* ε4 in ALS, we leveraged multiomic and longitudinal clinical data from the Answer ALS cohort. We tested whether *APOE* ε4 status modifies clinical trajectories and shapes iPSC-derived MN molecular profiles. We also asked whether *APOE* ε4 defines a distinct systemic plasma proteomic endophenotype (Fig. 1a). The Answer ALS cohort included 934 participants across 99 non-impaired controls, 774 ALS and 61 non-ALS MND cases. Controls and ALS cases were similar in age (55 ± 14 vs 55 ± 11 years), whereas the non-ALS MND group was older (62 ± 12 years). Participants were largely White across groups, and *APOE* genotypes were dominated by *APOE* ε3/ε3, with *APOE* ε4 carriage present in around 23–27% of participants depending on subgroup and *APOE* ε4/ε4 remaining rare (Supplementary Table 1).

**Figure 1.**
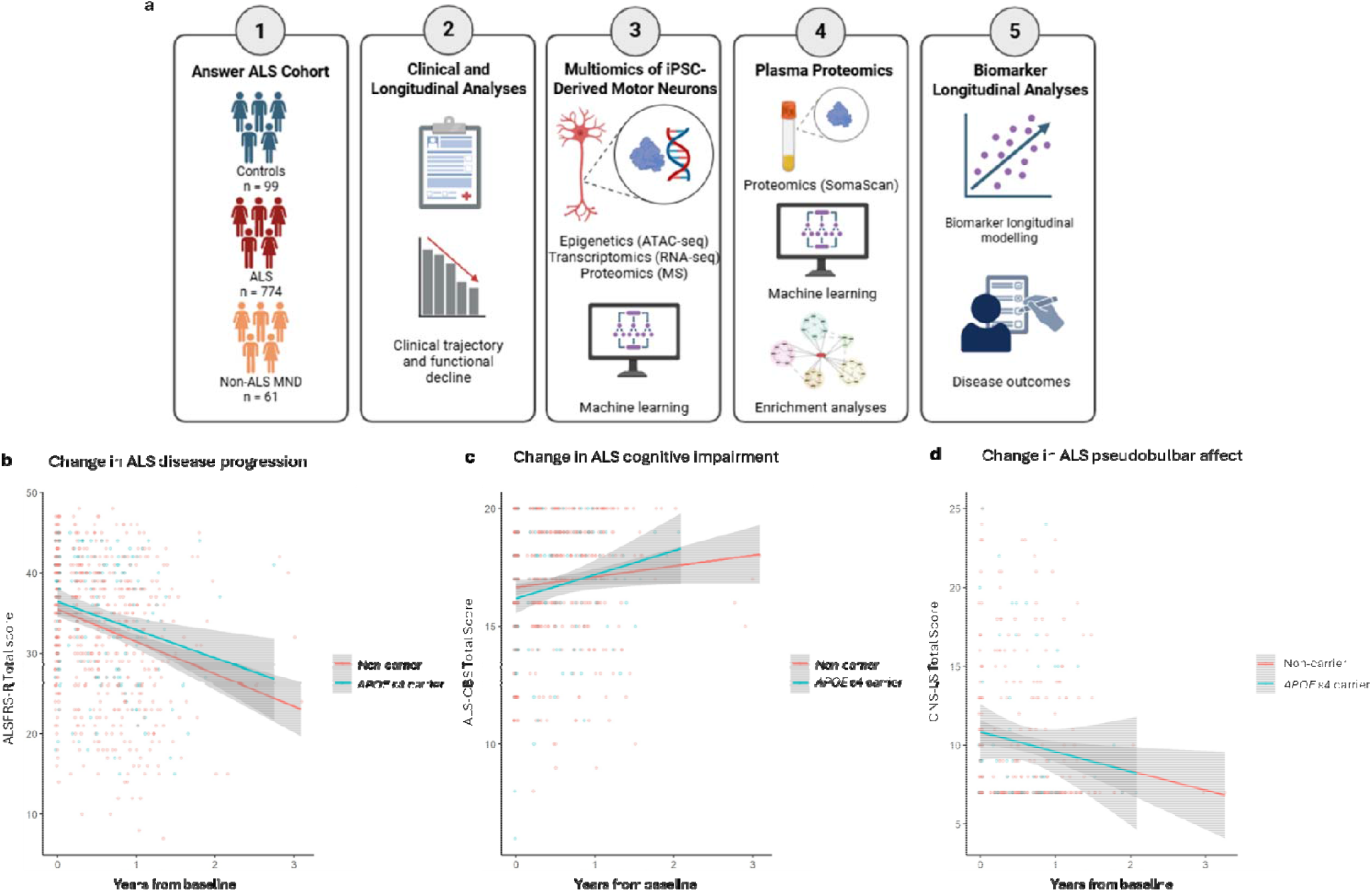
Study summary and *APOE* ε4 carriage is not associated with ALS clinical phenotype or longitudinal progression. (**a**) Summary of the current study. (**b-d**) Longitudinal trajectories of (**b**) ALS Functional Rating Scale-Revised (ALSFRS-R), (**c**) ALS Cognitive Behavioral Screen (ALS-CBS), and (**d**) Center for Neurologic Study-Lability Scale (CNS-LS) show no differences at baseline or over time between *APOE* ε4 carriers and non-carriers.

### *APOE* ε4 does not alter motor, cognitive, or neuropsychiatric trajectories in ALS

To assess whether *APOE* ε4 carriage influences the clinical course of ALS, we modelled longitudinal ALSFRS-R scores using linear mixed-effects models. Models included random intercepts and slopes to account for individual heterogeneity, and fixed effects for age, sex, time from baseline, *APOE* ε4 carrier status, and their interaction. There were no differences between *APOE* ε4 carriers and non-carriers on ALSFRS-R score at baseline (β = 0.80, SE = 1.07, t = 0.75, p = 0.45) and rate of functional decline was the same across both groups (*APOE* ε4 x time: β = 0.69, SE = 0.94, t = 0.74; Fig. 1b), with an average annual decrease of 5.8 ALSFRS-R points in carriers versus 6.5 in non-carriers. Cognitive performance, as measured by the ALS-CBS, similarly showed no *APOE* ε4-associated effect either at baseline (β = -0.23, SE = 0.43, t = -0.54) or over time (*APOE* ε4 x time: β = -0.06, SE = 0.48, t = -0.12; Fig 1c). Pseudobulbar affect, measured by CNS-LS total score, were similarly unaffected at baseline (β = 0.85, SE = 1.01, t = 0.85, p = 0.40). Longitudinal trajectories of CNS-LS in both *APOE* ε4 carriers and non-carriers remained flat (non-carriers: β = -0.04, SE = 0.49, t = -0.08; *APOE* ε4 x time: β = -1.04, SE = 1.11, t = -0.94; Fig. 1d). Together, these findings reinforce prior observations that *APOE* ε4 does not produce a clinically distinct ALS phenotype, and confirm that motor, cognitive, and affective decline are unaffected by *APOE* ε4 status.

### *APOE* ε4 does not affect the molecular profile of ALS motor neurons

Given the absence of APOE ε4-associated differences at the clinical level, we next asked whether *APOE* ε4 exerts functional effects at the cellular level by performing multiomic profiling of patient-derived iMNs. Chromatin accessibility, measured by ATAC-seq, revealed no differentially accessible regions associated with *APOE* ε4 status (Fig. 2a; Supplementary Table 2). Bulk RNA-sequencing similarly showed no substantial significant genotype-associated transcriptional changes. There were two genes (FDR < 0.05) that were differentially expressed, a previously uncharacterized gene (ENSG00000271254) and an antisense transcript overlapping CTSA (ENSG00000271984), the functional significance of which remains unclear (Fig. 2b; Supplementary Table 3). Label free mass spectrometry of the iMN proteome likewise revealed no differentially abundant proteins between *APOE* ε4 carriers and non-carriers (Fig. 2c; Supplementary Table 4). To determine whether *APOE* ε4 genotype effects might still be detectable in each omic dataset via higher-order, nonlinear patterns, we performed mutual information-based feature selection within each omic layer and trained random forest classifiers. None of the resulting models achieved strong discriminatory performance, with classification accuracy remaining modest and only slightly above random expectation (Table 1; Fig. 2d). These results suggest that *APOE* ε4 does not lead to detectable molecular changes in iMNs across chromatin accessibility, transcription, or protein abundance. While this reinforces the lack of overt clinical divergence, it also underscores that any *APOE* ε4-associated biological effect is unlikely to be neuron specific.

**Table 1.** Performance metrics of random forest classifiers on iMN individual omic datasets.

|  | **Sensitivity** | **Specificity** | **PPV** | **NPV** | **AUC** |
| --- | --- | --- | --- | --- | --- |
| ATAC-seq | 0.40 | 0.71 | 0.31 | 0.78 | 0.63 |
| RNA-seq | 0.33 | 0.80 | 0.37 | 0.77 | 0.55 |
| Mass spectrometry | 0.18 | 0.92 | 0.45 | 0.77 | 0.59 |

Abbreviations. AUC: area under the curve; NPV: negative predictive value; PPV: positive predictive value.

**Figure 2.**
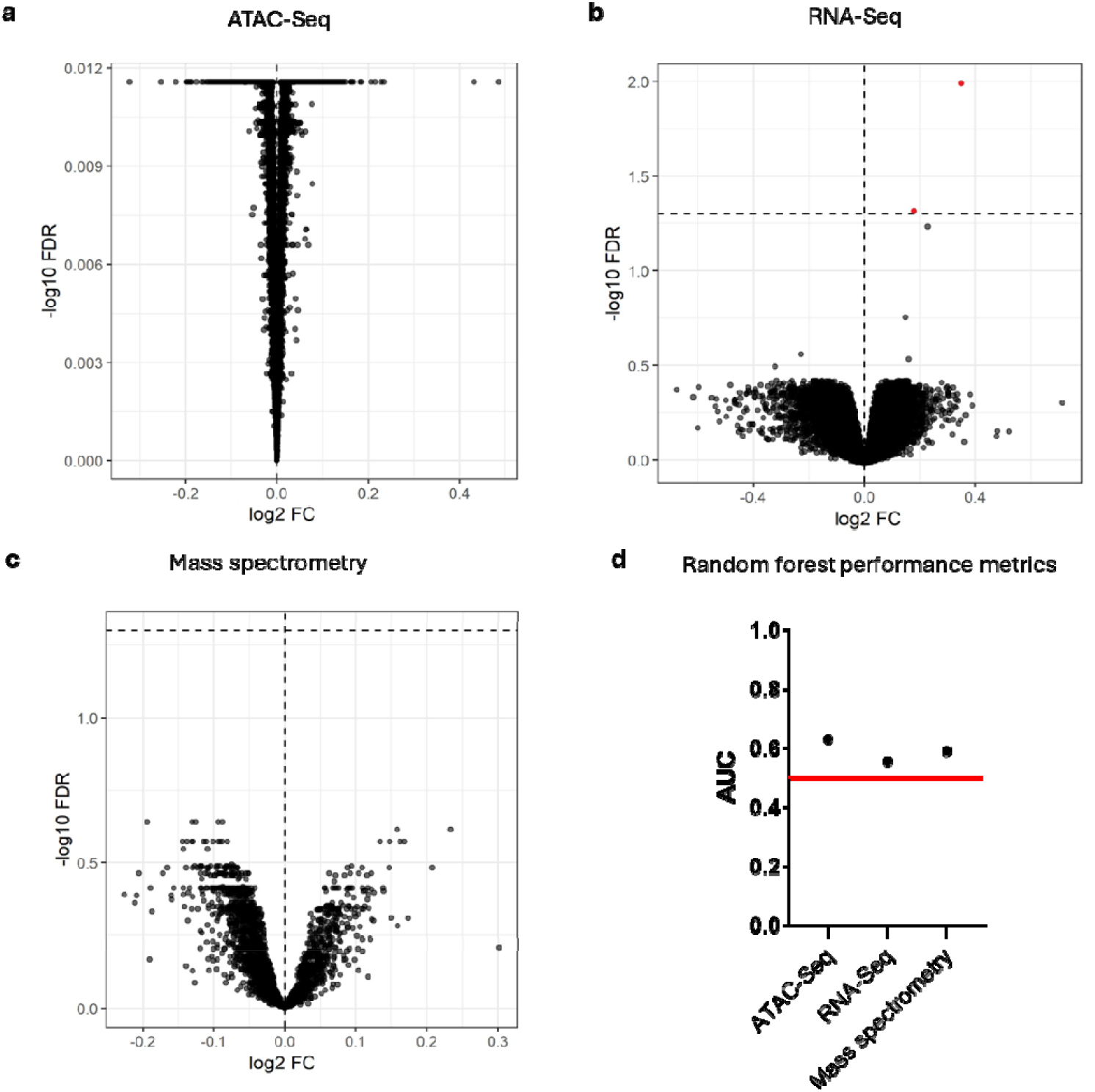
Multi-omic profiling of ALS patient induced pluripote ntstem cellderived motorneurons (iMNs)reveals no *APOE* ε4-associated differences. **(a-c)** Volcano plots showing differential (**a**) chromatin accessibility (ATAC-seq), (**b**) gene expression (RNA-seq), and (**c**) protein abundance (mass spectrometry) in iMNs from ALS patients who are *APOE* ε4 carriers versus non-carriers, plotted as log_2_ fold change versus -log_10_ adjusted *p*-value (false discovery rate; FDR). (**d**) Performance metrics of random forest classifiers indicate poor predictive power of chromatin accessibility, transcriptomic, and proteomic features for *APOE* ε4 status in ALS iMNs. The red line indicates chance performance (AUC = 0.50).

### APOE ε4 defines a systemic, immune-enriched plasma proteomic signature in ALS

Previous research from our group has showed that *APOE* ε4 has an immunomodulatory function in the context of neurodegenerative diseases, including ALS (42, 43). Given the lack of *APOE* ε4-associated differences in patient-derived iMNs and the lack of immune cells in these cultures, we next analysed plasma proteomics to capture potential *APOE* ε4 effects not observable in iMNs. Using mutual information-based feature selection, we identified a 44-protein plasma signature associated with *APOE* ε4 carrier status (Supplementary Table 2). Random forest classifiers trained on these proteins demonstrated robust performance, with high classification accuracy in both ALS (AUC = 0.98) and non-ALS MND (AUC = 0.86) participants (Table 2; Fig. 3a). Protein-protein interaction analysis revealed extensive network connectivity among the 44 proteins (Fig. 3b). Pathway enrichment analyses in KEGG and PANTHER showed the strongest overrepresentation for immune-related pathways, including the proteasome and FOXO signalling (Fig. 3c-i). Developmental programs such as meiosis were also implicated, likely due to shared involvement in DNA and RNA processing (Fig. 3c). Enrichment was also observed for cancer-related pathways, potentially reflecting underlying immune activation and proliferative signalling (Fig. 3h). In addition, the negative regulation of apoptosis pathway was enriched suggesting altered cell survival dynamics in *APOE* ε4 carriers (Fig. 3i).

**Table 2.**
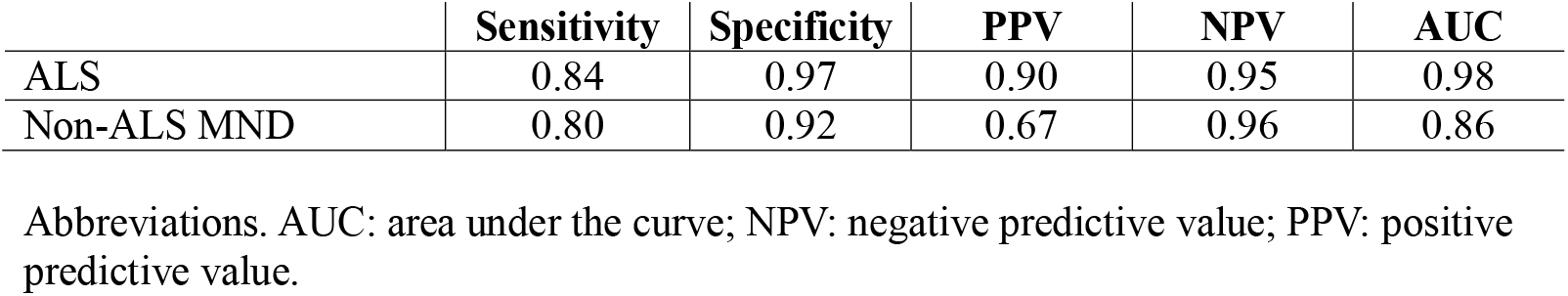
Performance metrics of random forest classifiers on plasma proteomic data.

|  | **Sensitivity** | **Specificity** | **PPV** | **NPV** | **AUC** |
| --- | --- | --- | --- | --- | --- |
| ALS | 0.84 | 0.97 | 0.90 | 0.95 | 0.98 |
| Non-ALS MND | 0.80 | 0.92 | 0.67 | 0.96 | 0.86 |

Abbreviations. AUC: area under the curve; NPV: negative predictive value; PPV: positive predictive value.

**Figure 3.**
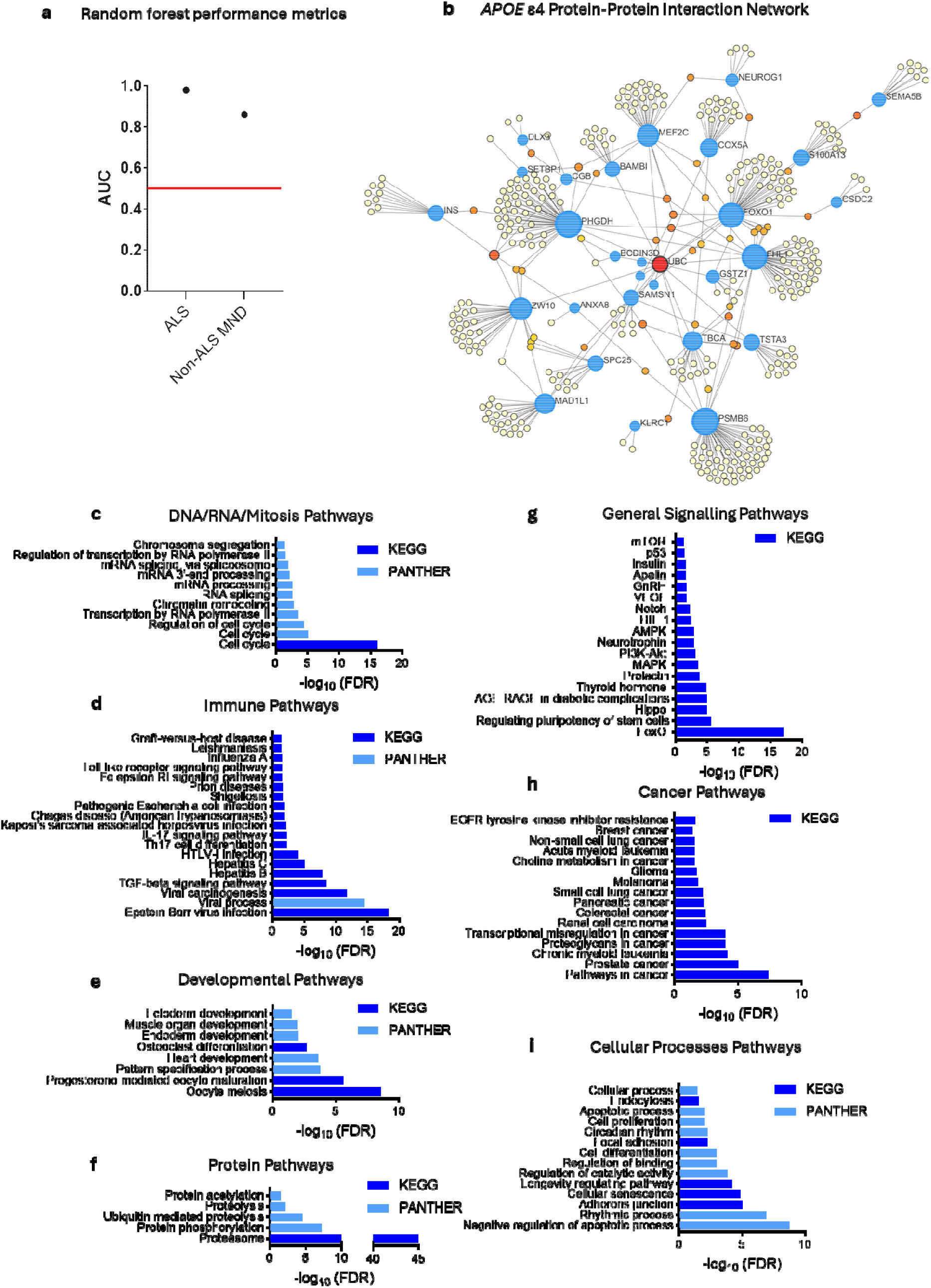
Plasma proteomic profiling reveals an *APOE* ε4-associated immune and inflammatory signature in ALS. (**a**) Performance of random forest classifiers demonstrates strong discrimination of *APOE* ε4 status using plasma proteomic features in ALS and non-ALS motor neuron disease (MND) participants; the red line indicates chance performance (AUC = 0.50). (**b**) Protein-protein interaction network of *APOE* ε4 plasma proteins, highlighting extensive interconnectivity. Blue nodes represent identified *APOE* ε4 proteins; red, orange, and yellow nodes indicate connected imputed downstream proteins, with colour denoting the degree of interconnection (red > orange > yellow). (**c-i**) Functional enrichment analysis of *APOE* ε4 plasma proteins (FDR < 0.05) showing over-representation of (**c**) DNA/RNA/mitotic processes, (**d**) immune pathways, (**e**) developmental processes, (**f**) protein-related pathways, (**g**) general signalling, (**h**) cancer-related pathways, and (**i**) cellular processes.

To explore the therapeutic relevance of this *APOE* ε4 plasma proteomic signature, we performed a protein-drug interaction analysis using the 44-protein set. This analysis revealed enrichment for several pharmacological classes, including carbonic anhydrase inhibitors (acetazolamide, benzthiazide, cyclothiazide, diclofenamide, ethoxzolamide, hydroflumethiazide, and methazolamide), antiallergic agents (amlexanox, olopatadine), nutritional supplements (calcium, ellagic acid, myristic acid, NADH, and zinc), and antidiabetic compounds (glisopexide, insulin, and IF402) (Table 3). These drugs converge on pathways involved in immune regulation, ion homeostasis, and muscle excitability, suggesting potential avenues for therapeutic modulation of *APOE* ε4-associated systemic dysfunction in ALS.

**Table 3.**
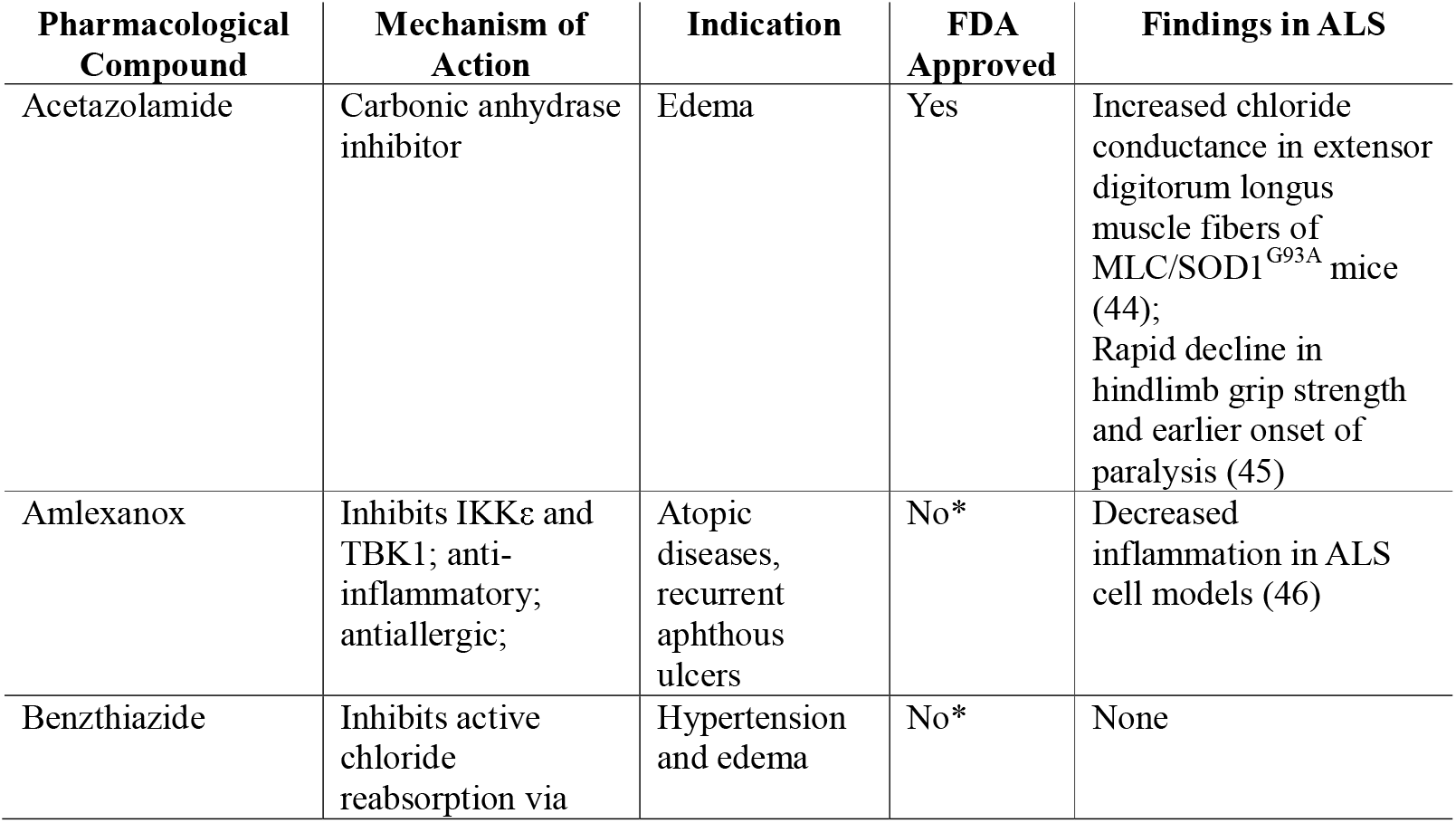
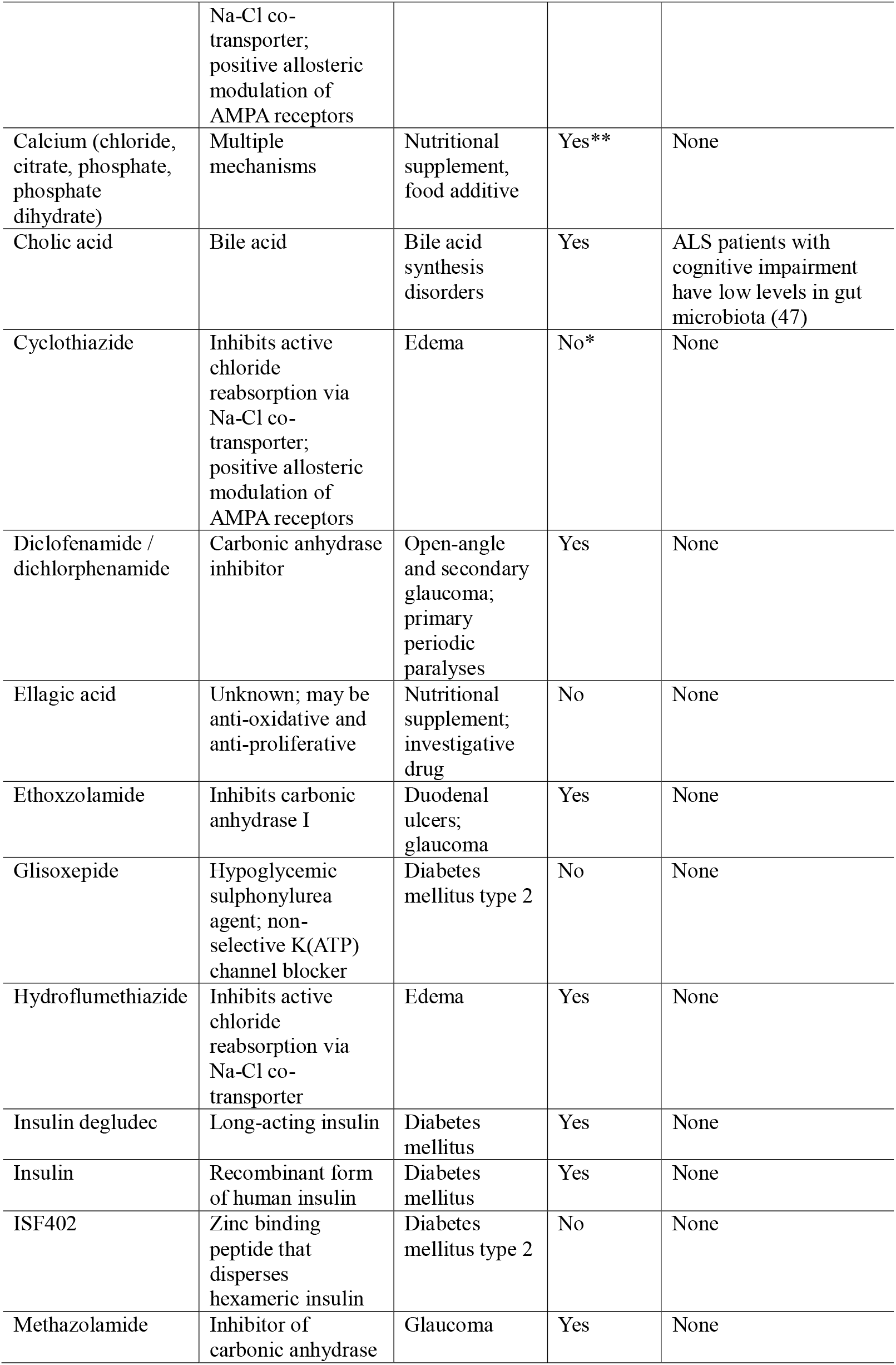
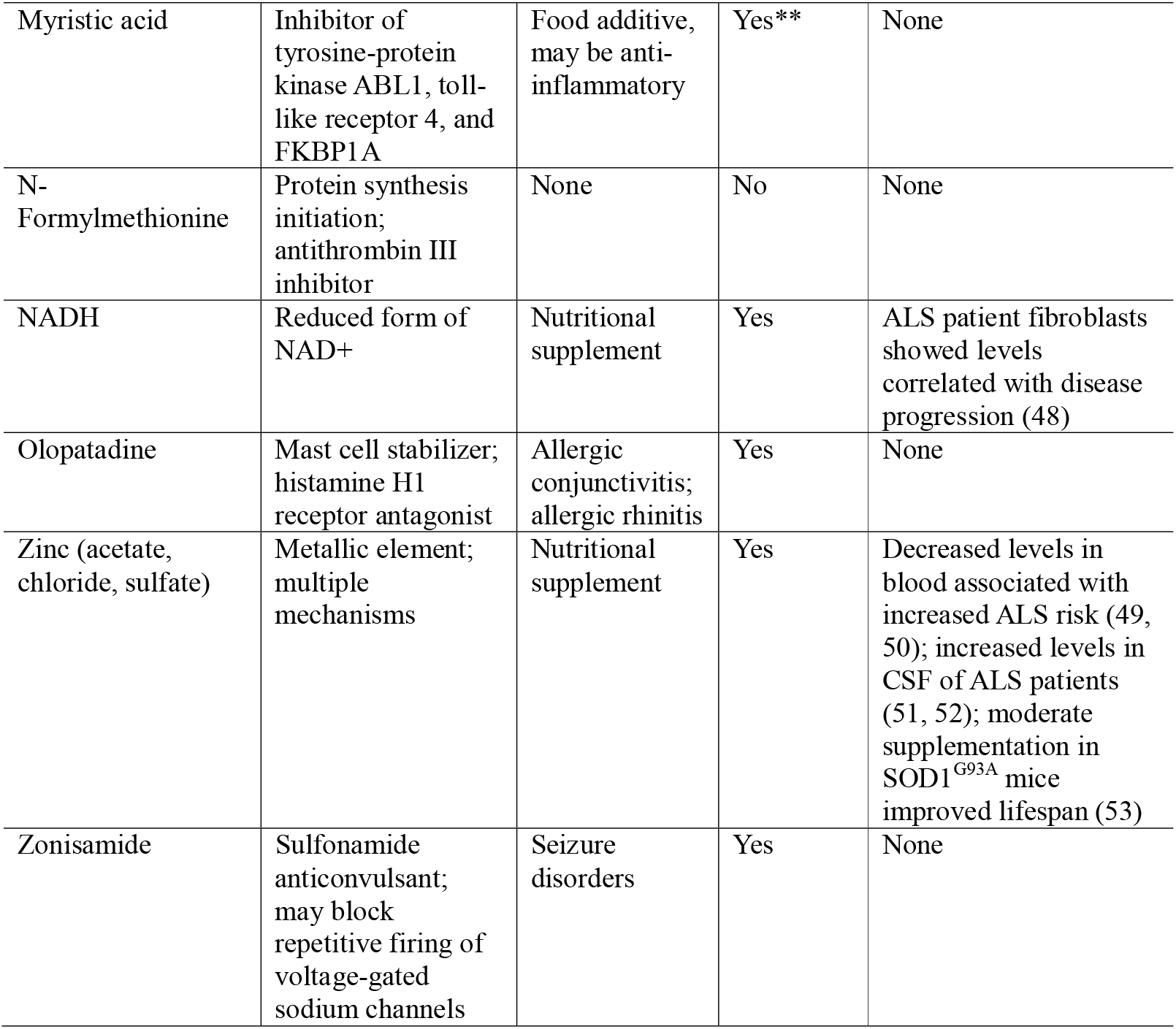
Pharmacological compounds interacting with *APOE* ε4 plasma proteins.

### *APOE* ε4 defines a stable, persistent plasma proteomic phenotype in ALS

To determine whether the *APOE* ε4 plasma proteomic signature reflects a transient or persistent systemic state, we evaluated its temporal stability across up to five longitudinal study visits. Effect sizes for *APOE* ε4 status on each of the 44 plasma proteins were estimated at each timepoint. These effects were highly concordant across visits, with pairwise correlations consistently exceeding r = 0.93 and near-perfect alignment between the first two timepoints (r = 0.99), indicating a highly stable APOE ε4 signal over time (Fig. 4a). Per-protein slopes of the *APOE* ε4 effect were centered near zero, and the mean absolute *APOE* ε4 effect across proteins remained consistent across all visits, with no evidence of temporal amplification or attenuation (Fig. 4b). To summarize this plasma protein pattern in a low-dimensional space, we derived an *APOE* ε4-associated axis by performing PCA on baseline protein concentrations in ALS participants. When longitudinal samples were projected onto this axis, *APOE* ε4 carriers consistently exhibited significantly lower PC1 scores than non-carriers at baseline (β = -2.99, SE = 0.26, t = -11.51). This separation remained stable throughout follow-up, with no significant effect of time (β = -0.06 units/year, SE = 0.10, t = - 0.63) and no evidence of a time-by-genotype interaction (β = 0.05 units/year, SE = 0.22, t = 0.23). To visualise individual-level dynamics of the *APOE* ε4 plasma proteins, we examined longitudinal trajectories of the baseline-derived PC1 score across study visits. *APOE* ε4 carriers and non-carriers remained stably separated over time, with no evidence of convergence or divergence at the group level (Fig. 4c). These findings demonstrate that the *APOE* ε4 systemic proteomic profile is not only robust, but temporally persistent, reflecting a stable endophenotype that precedes and accompanies disease progression.

**Figure 4.**
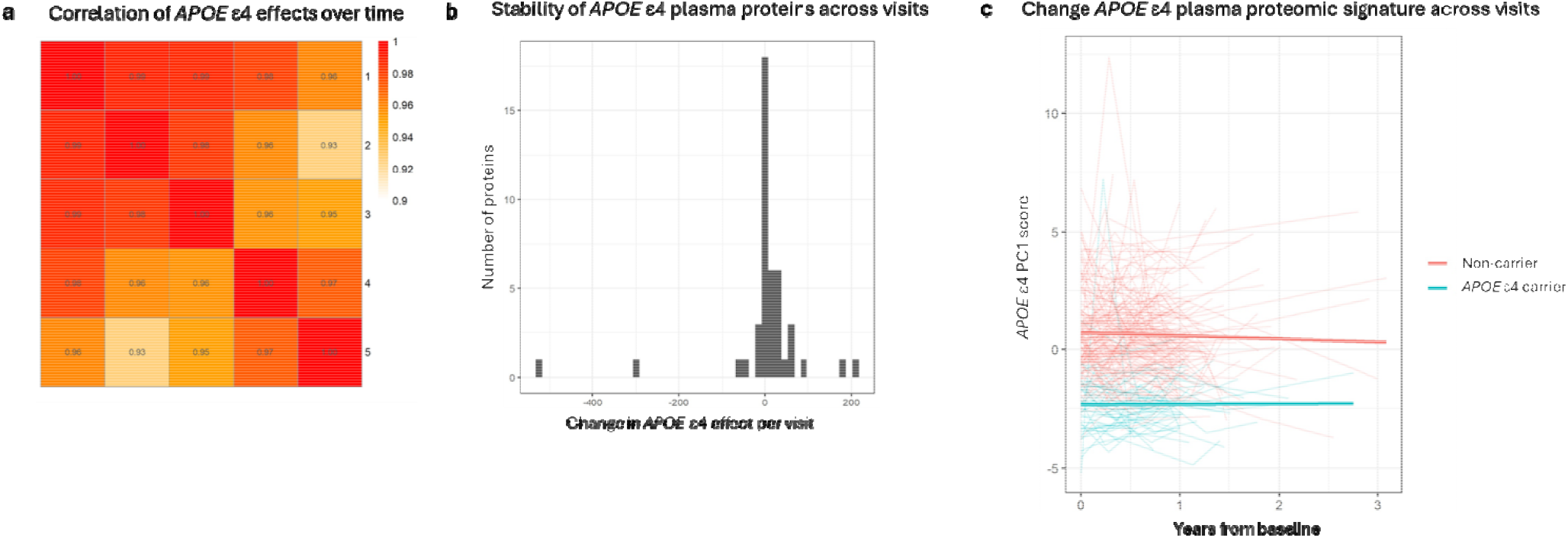
Temporal stability of the *APOE* ε4 plasma proteomic signature in ALS. (**a**) Correlation plot of effect sizes for *APOE* ε4 status on each of the 44 plasma proteins across visits one and two. (**b**) Histogram of the slopes of per-protein slopes of the *APOE* ε4 effect. (**c**) Longitudinal trajectories of individual participants projected onto the baseline *APOE* ε4 principal component 1 (PC1) axis. Solid lines represent group means ± standard error; individual trajectories are shown as semi-transparent lines.

To assess the clinical relevance of *APOE* ε4 plasma proteins, we first examined their relationship with longitudinal measures of ALS disease burden. Among the 44 candidate proteins, none showed significant associations with longitudinal change in cognitive function (CBS-ALS; Fig. 5a) or pseudobulbar affect (CNS-LS; Fig. 5b) after correction for multiple testing. In contrast, three proteins, PHGDH (O43175), TBCA (O75347), and NFL (P07196), demonstrated significant associations between their longitudinal slopes and ALSFRS-R decline (FDR < 0.05; Fig. 5c). For each, faster clinical deterioration (more negative ALSFRS-R slope) was associated with a more positive protein slope over time, suggesting preferential upregulation in rapidly progressing patients (Fig. 5d). To further highlight this, we derived a composite plasma biomarker score by extracting PC1 from the slopes of the three proteins. The plasma biomarker score was significantly correlated with ALSFRS-R slope (r = 0.28, 95% CI 0.15-0.40, p = 2.5 x 10^-5^) and remained strongly predictive in regression models (β = 0.998 ± 0.235 points/year per unit of PC1, p = 3.2 x 10^-5^), with no evidence of an effect of *APOE* ε4 carrier status on progression (p = 0.66) (Fig. 5e). It also significantly discriminated fast versus slow ALS progressors. In tertile-based models, each unit increase in PC1 was associated with increased odds of fast progression (OR = 1.59, 95% CI 1.19-2.13, p = 0.0017; Fig. 5f) and this effect remained robust when classifying participants by median ALSFRS-R slope (OR = 1.43, 95% CI 1.14-1.79, p = 0.0018). *APOE* ε4 status was not associated with progression group under either stratification (OR = 1.04, 95% CI 0.45-2.38, p = 0.93). Bootstrap analyses confirmed the robustness of the association, with the estimated effect direction preserved in 100% of resampled datasets and the odds ratio exceeding 1 in 99.95% of iterations. Permutation testing further supported the non-randomness of the association (permutation p = 0; logit-scale *p* = 0.001). Across bootstraps, NFL and PHGDH were reselected in 70% and 67% of iterations, respectively, while TBCA was included in 44%, indicating that the signal is driven by a stable proteomic core rather than sampling noise.

**Figure 5.**
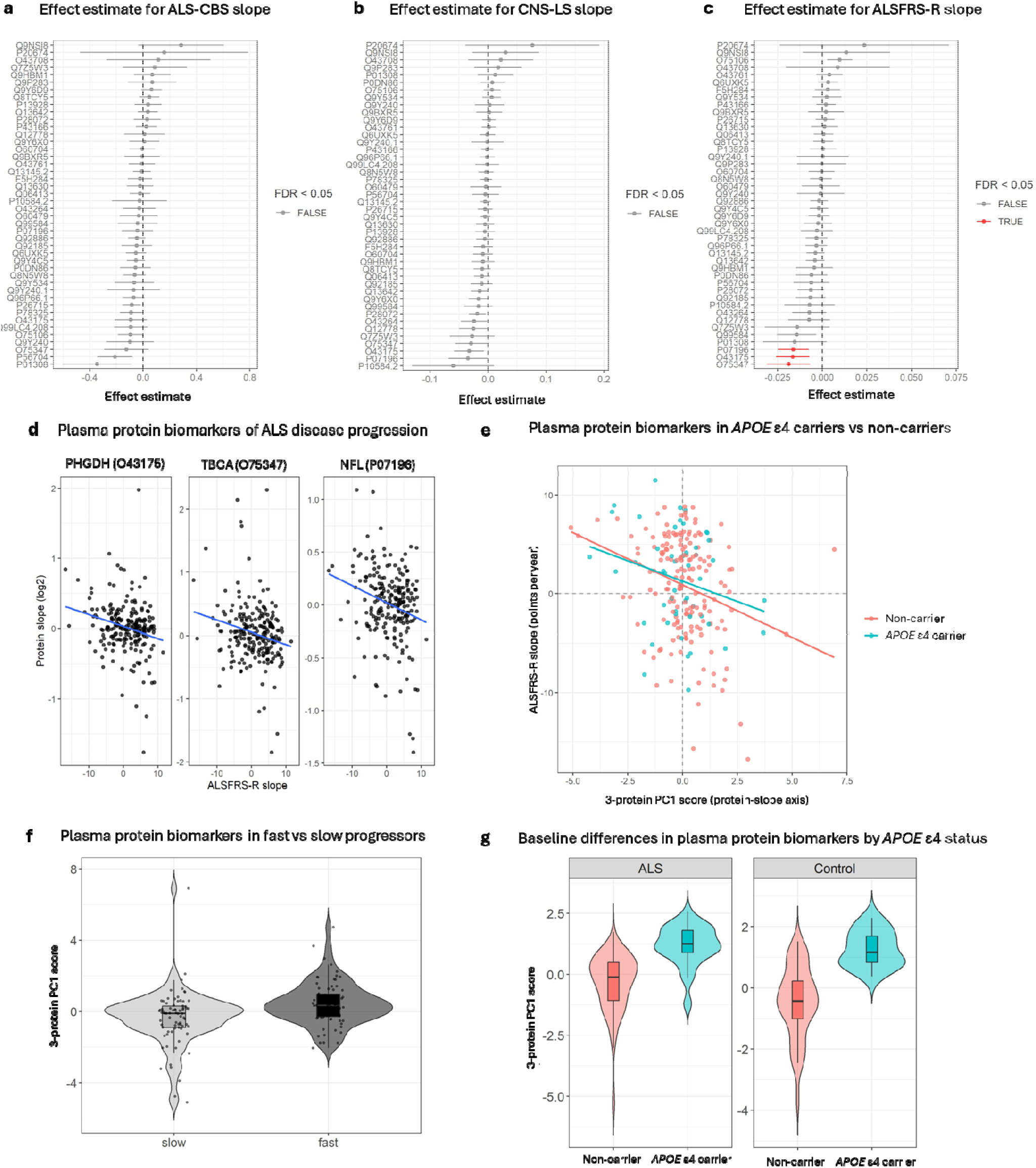
Association of *APOE* ε4 plasma proteins with ALS disease progression and baseline genotype effects. (**a-c**) Forest plots showing associations between longitudinal protein slopes and longitudinal change in (**a**) cognitive function (CBS-ALS), (**b**) pseudobulbar affect (CNS-LS), and (**c**) ALS Functional Rating Scale–Revised (ALSFRS-R). Points indicate regression coefficients and horizontal bars denote 95% confidence intervals. Proteins passing false discovery rate (FDR) correction are highlighted in red. (**d**) Scatter plots showing the relationship between longitudinal protein slopes and ALSFRS-R slope for the three significant proteins (PHGDH, TBCA, and NFL). Each point represents an individual participant; solid lines indicate fitted linear regression trends. More rapid clinical decline (more negative ALSFRS-R slope) is associated with increasing protein abundance over time. (**e**) Relationship between ALSFRS-R slope and the composite plasma biomarker score derived as principal component 1 (PC1) of the three protein slopes. Lines indicate fitted regression models stratified by *APOE* ε4 carrier status. (**f**) Violin plots showing distribution of the plasma biomarker score in fast versus slow ALS progressors, defined by median ALSFRS-R slope. Boxplots denote median and interquartile range. (**g**) Distribution of the plasma biomarker score stratified by *APOE* ε4 carrier status in ALS patients and neurologically healthy controls at baseline.

We next assessed whether APOE ε4 status was associated with baseline differences in the plasma biomarker score. Notably, the three plasma biomarker proteins were initially identified as *APOE* ε4-associated features, suggesting that carriers exhibit a distinct proteomic baseline that may shape disease expression independently of progression rate. In healthy controls, the plasma biomarker score was significantly higher in *APOE* ε4 carriers (β = 1.71 ± 0.31 SD units, *p* = 8.7 x 10^-7^, R^2^ = 0.35), with a similarly large effect in ALS patients at baseline (β = 1.51 ± 0.19, *p* = 4.4 × 10^-14^, R^2^ = 0.29; Fig. 5g). Longitudinal mixed-effects models further demonstrated that this *APOE* ε4-associated shift was time-invariant (β = 1.77 SD units, t = 10.5), with no significant main effect of time (*p* = 0.53) or time-by-genotype interaction (p = 0.82). Together, these findings support a model in which *APOE* ε4 defines a stable, systemic baseline that is detectable even in neurologically healthy individuals.

## Discussion

In this study, we addressed the longstanding debate over *APOE* ε4 in ALS by leveraging the data from the large-scale Answer ALS cohort. Consistent with prior reports (3, 4, 5, 7, 8, 9, 10, 14, 54), *APOE* ε4 genotype was not associated with clinical onset, progression, or phenotypic variation in ALS. *APOE* ε4 status also had no detectable impact on chromatin, transcriptional, or proteomic profiles in patient-derived iMNs, reinforcing the prevailing view that it does not drive overt neuronal or clinical divergence. However, this interpretation conflates the absence of a genotype-phenotype association with biological irrelevance. Our findings challenge that assumption, revealing a robust and longitudinally stable plasma proteomic signature associated with *APOE* ε4 and enriched for immune and inflammatory pathways. While *APOE* ε4 itself did not change the relationship between these proteins and ALS progression, several proteins within the signature were independently associated with faster ALSFRS-R decline and higher in *APOE* ε4 carriers. This suggests that *APOE* ε4 configures a peripheral immune context that persists over time and may influence disease expression through non-cell-autonomous mechanisms. Therefore, *APOE* ε4 is biologically relevant in ALS despite not leading to changes in clinical trajectory. Together, these findings demonstrate that the absence of an overt clinical or neuronal phenotype does not imply biological neutrality. Instead, they position *APOE* ε4 as a systemic immune modulator that shapes the molecular landscape of ALS without altering genotype specific clinical trajectories.

A key insight from our study is that *APOE* ε4’s effects are context-dependent, emerging in immune-rich environments but not in isolated neurons. In patient-derived iMNs, we observed no *APOE* ε4-associated differences in chromatin accessibility, gene expression, or protein abundance, likely reflecting the absence of glial and immune cells in these neuron-only cultures. *APOE* is predominantly expressed by astrocytes and microglia in the CNS and plays critical roles in lipid transport and injury response (55). The absence of a cell-autonomous *APOE* ε4 signal therefore does not argue against biological relevance but rather suggests that *APOE* ε4 exerts its effects through interactions with immune cells. In support of this, plasma proteomics revealed a robust *APOE* ε4-associated immune signature, implicating systemic pathways not captured by neuronal assays. This aligns with our previous work showing *APOE* ε4 acts as an immune modulator (42, 43) and with other studies showing that *APOE* ε4 homozygosity is associated with inflammatory signatures and disease-associated microglial states in ALS (13). Mechanistically, several pathways enriched in the *APOE* ε4 plasma proteome provide further support for this idea. We found substantial significant enrichment for diverse pro-inflammatory, viral, and immune pathways. Additionally, we found enrichment for FOXO signalling and the proteosome. FOXO plays a central role in both innate and adaptive immunity, regulating the development and differentiation of dendritic cells, B cells, and multiple T cell subsets (56, 57, 58). Similarly, the proteasome pathway governs inflammatory signalling, protein turnover, and immune cell differentiation (59). The enrichment of these pathways in *APOE* ε4 carriers underscores a broader role for APOE ε4 in modulating systemic immune function and inflammatory signalling to influence ALS pathophysiology. Importantly, the lack of an *APOE* ε4-associated difference in ALSFRS-R trajectory does not diminish the relevance of these findings. Clinical scales capture late, integrative disease outcomes, whereas molecular endophenotypes provide insight into upstream biological processes that may be therapeutically actionable. Identifying stable, genotype-associated immune signatures therefore informs disease mechanism and target discovery, even in the absence of genotype-specific differences in overt clinical progression. This distinction is central to precision medicine approaches, where biological stratification can reveal vulnerabilities not apparent at the phenotypic level.

From a translational perspective, the *APOE* ε4-associated plasma proteomic signature may offer new therapeutic insights, particularly for stratified intervention in *APOE* ε4-positive ALS patients. Drug enrichment analysis of this signature identified several pharmacological classes converging on inflammation, ion channel regulation, and oxidative stress, pathways implicated in ALS pathophysiology. One notable class was carbonic anhydrase inhibitors, including acetazolamide, dichlorphenamide, ethoxzolamide, and methazolamide. These agents are established treatments for hypokalemic periodic paralysis, where they modulate calcium-activated potassium channels in muscle fibers to stabilize membrane excitability (60, 61). Dichlorphenamide improves episodic weakness and is well tolerated in randomized trials (62, 63, 64) and acetazolamide remains a first-line treatment (65). While periodic paralysis and ALS are fundamentally distinct, non-degenerative versus neurodegenerative, the emergence of this drug class in our analysis raises the possibility that targeting excitability or early metabolic stress responses could be beneficial during the earlier stages of ALS. Zonisamide, another enriched compound, is a T-type calcium and sodium channel blocker with weak carbonic anhydrase inhibitory activity. It has demonstrated neuroprotective effects in PD models by reducing oxidative stress, apoptosis, and neuroinflammation (66, 67, 68, 69). We also identified amlexanox, an inhibitor of TBK1/IKKε, which is of particular interest given that TBK1 is a known ALS risk gene (70). While amlexanox decreased neuroinflammation in animal models of epilepsy (71), it showed toxicity in iMNs from a Charcot-Marie-Tooth patient (72), underscoring the importance of functional validation in disease-relevant contexts. Zinc-related compounds also emerged from our analysis. Zinc dysregulation has been variably implicated in ALS. CSF levels are elevated in patients (51, 52), early life zinc exposure has been associated with increased ALS risk (73), and supplementation shows dose-dependent effects in SOD1^G93A^ mouse models, ranging from lifespan extension to lethal anemia depending on dosage (53, 74). Blood zinc levels have also been inversely correlated with ALS risk in population studies (49, 50). These conflicting findings underscore the complexity of zinc biology but also highlight the inflammatory and oxidative stress consequences of deficiency and potential for therapeutic modulation in ALS (75, 76, 77). Together, these findings nominate several candidate compounds that converge on key processes implicated in *APOE* ε4-associated ALS. While preliminary, they offer tractable precision therapeutic strategies for preclinical investigation in *APOE* ε4 carriers with ALS. Future research would benefit from testing these compounds in more complex human disease-specific models, such as iPSC-derived organoids, that can accurately model *APOE* ε4’s effects (78, 79).

Our analyses identified PHGDH, TBCA, and NFL as components of an *APOE* ε4- associated plasma proteomic signature with relevance to ALS disease progression, broadly. PHGDH, the rate-limiting enzyme of *de novo* serine biosynthesis, links metabolic state to redox and inflammatory control. While PHGDH activity supports cellular resilience by maintaining serine and glutathione availability, protecting against senescence and oxidative damage (80), its sustained upregulation may also reflect maladaptive stress responses. In the CNS, PHGDH is highly expressed in glial populations (81) and promotes a pro-inflammatory microglial phenotype characterized by increased NO, IL-6, and TNF production (82). PHGDH also contributes to skeletal muscle biomass generation (83, 84), suggesting that it may be a biomarker that is reflective of broader dysfunctions in ALS spanning immune, metabolic, and neuromuscular systems. TBCA, a tubulin-specific chaperone required for β-tubulin folding and microtubule assembly (85), implicates cytoskeletal vulnerability as another biomarker for ALS progression. Disruption of tubulin homeostasis is a recognized feature across neurodegenerative disorders, with neurofibrillary tangle-like β-tubulin structures reported in ALS, AD, Guam ALS/parkinsonism-dementia complex, and Down syndrome (86). NFL provides a complementary and clinically anchored signal. Although NFL lacks disease specificity and is elevated across a wide range of neurological and inflammatory conditions (87), its prognostic utility in ALS is well established. Serum NFL levels correlate with axonal injury, predict more rapid functional decline, and differ systematically across ALS genetic subtypes, with the highest levels observed in C9orf72-associated disease (88, 89, 90). Importantly, the elevation of PHGDH, TBCA, and NFL in *APOE* ε4 carriers was evident early in disease, supporting a model in which *APOE* ε4 establishes a pre-existing biological vulnerability rather than driving genotype-specific clinical trajectories. This interpretation is consistent with cross-disease proteomic analyses demonstrating shared immune-related alterations in *APOE* ε4 carriers during preclinical or prodromal neurodegeneration (42, 43, 79, 91). Collectively, these findings position *APOE* ε4-associated plasma proteins not merely as correlates of disease activity, but as indicators of an underlying immune-metabolic and structural state that may shape ALS pathogenesis and progression from the outset.

Our study has several limitations. First, our multiomic analyses were performed on iPSC-derived motor neurons in monoculture, lacking glia and peripheral immune cells, key mediators of *APOE* ε4 biology. This system restricts interpretation to neuron-intrinsic effects and likely omits critical neuroimmune interactions. However, this reductionist model also offers a strength. It enables direct testing of whether *APOE* ε4 alters neuronal epigenomic, transcriptomic, or proteomic states in isolation. The absence of *APOE* ε4-dependent effects in iMNs suggests that its pathogenicity in ALS is not cell-autonomous but instead requires interactions with the immune or glial environment. This interpretation aligns with the *APOE* ε4-associated immune signature observed in plasma, underscoring the need for future studies using multicellular systems that better reflect the disease context. Second, our study did not include cerebrospinal fluid or brain tissue from *APOE* ε4 carriers with ALS, limiting our ability to determine whether the plasma proteomic signature reflects CNS-specific processes. However, in previous work, we analyzed dorsolateral prefrontal cortex samples from *APOE* ε4 carriers with ALS and observed proteomic profiles broadly consistent with those seen in plasma (43) suggesting at least partial overlap between peripheral and CNS signatures. Future studies should extend this investigation to include spinal cord and motor cortex, the primary sites of neurodegeneration in ALS, to assess the extent to which the plasma *APOE* ε4 signature is mirrored within disease-relevant CNS regions. Finally, although the therapeutic implications we propose are based on bioinformatic enrichment analyses and prior pharmacological literature they remain speculative in the absence of functional validation. While compounds such as acetazolamide, amlexanox, and zonisamide have mechanistic plausibility, their efficacy in modifying *APOE* ε4-associated disease biology in ALS has yet to be demonstrated. Preclinical testing in relevant models will be essential to determine whether these compounds can attenuate the consequences of *APOE* ε4 in ALS. Furthermore, because *APOE* ε4 influences a range of interconnected biological processes, it is unlikely that a single therapeutic agent will fully mitigate its pathological effects. Instead, optimal therapeutic strategies may require a combination of interventions that simultaneously address multiple components of the *APOE* ε4-driven biological vulnerability to neurodegeneration.

## Conclusions

In a large longitudinal ALS cohort, *APOE* ε4 carriage did not stratify clinical trajectories and was not captured by patient iMN multiomics, consistent with a limited motor neuron-intrinsic *APOE* ε4 signal in ALS. Plasma proteomics, however, revealed a robust *APOE* ε4-associated protein signature that classified genotype with high accuracy across ALS and non-ALS motor neuron disease and was strongly enriched for immune and pro-inflammatory signalling. Critically, this systemic signature was highly stable over time, defining a persistent genotype-linked endophenotype rather than a transient disease state correlate. Within the *APOE* ε4 plasma proteomic signature, a distinct set of proteins showed reproducible associations with longitudinal functional decline and, when integrated as a composite, stratified individuals with more rapid versus slower trajectories. Notably, this progression-linked signal was already elevated at baseline in *APOE* ε4 carriers, including non-impaired controls, indicating a stable systemic endophenotype rather than a simple case-control effect. Collectively, these data show that an absent or weak *APOE* ε4 association with clinical phenotype does not imply biological neutrality. Instead, *APOE* ε4 is linked to a stable, immune-enriched peripheral endophenotype in ALS that partially overlaps with proteomic correlates of disease progression. By resolving an *APOE* ε4-linked plasma immune signature despite minimal clinical stratification, this study demonstrates the value of genotype-informed plasma proteomics for uncovering potential mechanistic functional consequences that are not evident from phenotype alone. Together, these findings provide a clear rationale for *APOE*-stratified biomarker development in ALS as an early step toward genotype-informed precision medicine. More broadly, they establish a generalizable template for testing whether other neurodegeneration-associated genetic variants define stable, peripherally measurable molecular endophenotypes with functional relevance to disease course.

## List of abbreviations

*ANG*: angiogenin
ALS: amyotrophic lateral sclerosis
ALS-CBS: ALS Cognitive Behavioral Screen
ALSFRS-R: ALS Functional Rating Scale-Revised
*APOE* ε4: apolipoprotein E ε4
AUC: area under the curve
*C9orf72*: chromosome 9 open reading frame 72
CNS-LS: Center for Neurologic Study-Lability scale
CSF: cerebrospinal fluid
FOXO: forkhead box O
*FUS*: FUS RNA binding protein
iPSC: induced pluripotent stem cell
iMNs: iPSC-derived motor neurons
KEGG: Kyoto Encyclopedia of Genes and Genomes
*MAPT*: microtubule-associated protein tau
MND: motor neuron disease
NFL: neurofilament light
NPV: negative predictive value
PANTHER: protein analysis through evolutionary relationships
PD: Parkinson’s disease
*PGRN*: progranulin
PHGDH: phosphoglycerate dehydrogenase
PPV: positive predictive value
*SETX*: senataxin
*SOD1*: superoxide dismutase 1
*TARDBP*: TAR DNA binding protein
*TBK1*: TANK-binding kinase 1
TBCA: tubulin folding cofactor A
TREM2: triggering receptor expressed on myeloid cells 2
*VAPB*: VAMP-associated protein B and C
*VCP*: valosin-containing protein.

## Declarations

## Ethics approval and consent to participate

Ethics approval was obtained by the respective contributing clinical sites in the Answer ALS Consortium. All participants provided informed consent to participate.

## Consent for publication

Not applicable.

## Availability of data and materials

The Answer ALS Consortium dataset (v3) was used and analysed in the current study. It is available through the Neuromine data portal: https://dataportal.answerals.org/. Researchers who wish to access this controlled dataset are required to register and submit a Data Use Agreement. Summary results generated in this manuscript are available in Supplementary Tables (S2-S5).

## Author contributions

A.S. and C.A.F. conceived the paper and designed the study; A.S., S.T., S.K., and C.A.F. performed the analyses; T.G.T. and J.D.R. provided key insights into the Answer ALS cohort. A.S. and C.A.F. made the figures and wrote the manuscript. All authors provided feedback on the manuscript and approved the final version.

## Acknowledgements

The authors are grateful to the Answer ALS Consortium, clinical cohort contributors, patients, donors, and families who helped to make this research possible. Data from the ALS TDI ARC Study and Answer ALS Data Portal (AALS-01184) were used in this work.

